# Resilience to cardiac aging in Greenland shark *Somniosus microcephalus*

**DOI:** 10.64898/2025.12.20.695706

**Authors:** Elena Chiavacci, Kirstine Fleng Steffensen, Pierre Delaroche, Emanuele Astoricchio, Amalie Bech Poulsen, Daniel Brayson, Fulvio Garibaldi, Luca Lanteri, Christian Pinali, Giovanni Roppo Valente, Federico Vignati, John Fleng Steffensen, Holly Shiels, Eva Terzibasi Tozzini, Alessandro Cellerino

## Abstract

The Greenland shark (*Somniosus microcephalus*), with a lifespan exceeding 400 years, represents a unique model for studying vertebrate longevity. Here, we characterize its cardiac aging profile and compare it with two other species: the deep-sea shark *Etmopterus spinax* and the short-lived teleost *Nothobranchius furzeri*. Histological analysis revealed extensive interstitial and perivascular fibrosis throughout the ventricular myocardium of *S. microcephalus*, affecting both compact and spongy layers of both sexes. This fibrotic pattern was absent in *E. spinax* and *N. furzeri*, suggesting it is a specific feature of *S. microcephalus*. We also observed extreme lipofuscin accumulation within cardiomyocytes of *S. microcephalus*, which correlates at the ultrastructural level with abundance of damaged mitochondria and the presence of strikingly enlarged lysosomes filled with electrondense material of likely mitochondrial origin. Additionally, in the myocardium of *S. microcephalus* we found abundant deposition of the oxidative stress marker 3-nitrotyrosine. Remarkably, despite showing multiple canonical markers of aging such as fibrosis, lipofuscin accumulation, and oxidative damage, S. microcephalus individuals appeared healthy and physiologically uncompromised at the time of capture. These findings suggest that *S. microcephalus* has evolved resilience to molecular and tissue-level aging hallmarks, supporting sustained cardiac function over centuries and offering new insights into the mechanisms of extreme vertebrate longevity.

## 1. Introduction

The Greenland shark (*Somniosus microcephalus*) is a giant (>5m) shark living in deep, cold water in high northern latitudes. It is characterized by slow growth rate (∼1cm/year) and exceptional longevity. Radiocarbon dating of eye lens nuclei revealed that sexual maturity is reached at ∼ 150 years and a maximum lifespans is at least 272 years, making it the slowest maturing, longest-living known vertebrate (Hansen 1963; Nielsen et al. 2016). *S. microcephalus* is a sluggish swimmer, with average cruising speed of ∼ 0.3m/s and ∼ 9 tail beats per minute, with possibly little capacity for sustained high-speed swimming (Compagno et al. 2005; Watanabe et al. 2012a; Nielsen et al. 2016). When corrected for body size, the Greenland shark exhibits the slowest sustained swim speed and tail-beat frequency recorded among known fishes (Watanabe et al. 2012b) matched by a low metabolism (Ste-Marie et al. 2022). Despite its remarkable longevity, knowledge of life history, ecology and physiology of this charismatic and elusive animal is very limited (MacNeil et al. 2012; Costantini et al. 2017; Herbert et al. 2017). Given the preeminent contribution of cardiovascular function to general mortality risk (Visseren et al. 2021; Vlachopoulos et al. 2010; Timmis et al. 2024; Townsend et al. 2022), a particularly interesting question is how heart function can be maintained for centuries in this species. In humans and most vertebrates, aging is characterized by apparent histological changes in the hearts, including cardiac fibrotic remodeling, loss of cardiomyocyte reserve, and cumulative oxidative damage, all of which contribute to the heart functional decline (Biernacka & Frangogiannis 2011; Lu et al. 2017; Fleg & Strait 2012; Yoneyama et al. 2016; Anversa & Nadal-Ginard 2002; Anversa & Nadal-Ginard 2002; Gazoti Debessa et al. 2001; Chen et al. 2022; Lesnefsky et al. 2016). Recently, spatial RNA sequencing in old mouse heart reveals a clear picture of the age-related changes in gene expression toward a fibrotic cardiac phenotype, while proteomics data showed an increase of reactive oxygen species (ROS) and a concomitant decline in antioxidant defenses which exacerbate the oxidative stress (Basilicata et al. n.d.). It remains unknown whether *S. microcephalus* is spared by such age-related cardiac remodeling and related functional consequences. Some characteristics of the Greenland shark’s heart and blood vessels have only recently begun to emerge. *S. microcephalus* has a two-chambered heart typical of Chondrichthyans, and consists of sinus venosus, atrium, ventricle and outflow tract arranged sequentially, pumping blood to the gills for oxygenation before circulating it through the body (Shadwick et al. 2018). *S. microcephalus* cardiocirculatory system may exhibit specific physiological and structural adaptations to its sluggish lifestyle that makes it resistant to age-dependent remodelling, thereby enabling its extreme longevity. An alternative possibility is that *S. microcephalus* maintains its cardiac function despite age-dependent remodelling thereby showing resilience.

In the present study, we set to discriminate between these two possibilities and investigated histological aging hallmarks in *S. microcephalus* heart specimens. We also analyzed hearts of two key comparison species: the deep-sea shark *Etmopterus spinax* (Linnaeus, 1758) and the short-lived African turquoise killifish (*Nothobranchius furzeri*). *E. spinax*, commonly known as the velvet belly lantern shark, distributed widely across the northeast Atlantic Ocean and the Mediterranean Sea (Compagno, L. 1984; Aranha et al. 2009). Both *E. spinax* and *S. microcephalus* derive from a common Squalomorph ancestor that lived ∼ 100 Mya and colonized deep-sea habitats (Straube et al. 2015), but *E. spinax* is of minute size (< 50 cm) and wild-capture records indicate maximum lifespans of ∼ 8 years for males and ∼ 11 for females (Gennari & Scacco 2007; Coelho & Erzini 2008), making it a valuable comparative model for distinguishing traits specifically associated with exceptional lifespan from general deep-sea adaptations. *N. furzeri* is the vertebrate with the shortest captive lifespan and is an emerging vertebrate model in aging research due to its compressed lifespan and expression of all the key hallmarks of mammalian aging (Cellerino et al. 2016; Di Cicco et al. 2011). Moreover, it serves as a key reference for cardiac aging in a poikilothermic aquatic vertebrate (Ahuja et al. 2019).

Lipofuscin is an autofluorescent, yellow-brown pigment composed primarily of cross-linked oxidized proteins, lipids, and metals. It accumulates progressively in postmitotic cells over time and is widely recognized as an hallmark of cellular aging (Terman & Brunk 1998; Terman & Brunk 2006; Brunk & Terman 2002; Malkoff & Strehler 1963; Kakimoto et al. 2019). In *N. furzeri* lipofuscin age-related accumulation was employed as an aging marker in the liver (Ng’oma et al. 2014; Terzibasi et al. 2009), brain (Terzibasi et al. 2008; Terzibasi Tozzini et al. 2013) and heart (Ahuja et al. 2019). Furthermore recently *N. furzeri* has also been proposed as a valuable model for studying cardiac aging (Ahuja et al. 2019; Ma et al. 2025). Furthermore, the heart of aged *N. furzeri* exhibits accumulation of 3-nitrotyrosine (3-NT), another well-established hallmark of aging due to oxidative and nitrosative stress. In rat models, 3-NT levels rise progressively with age, with elevated 3-NT levels linked to impaired SERCA2a function, mitochondrial destabilization, and endothelial dysfunction, even in the absence of overt cardiovascular disease (Knyushko et al. 2005), this is consistent with a gradual disruption in redox homeostasis during physiological cardiac aging. Similarly, NT-3 was found increased in aged *N. furzeri* healthy hearts, indicating 3-NT as reliable marker of cardiac aging (Heid et al. 2017). Thus we included the aged teleost model *N. furzeri* as a natural comparative model to the exceptionally long-lived Chondrichthyan *S. microcephalus.* The comparison of the three species *S. microcephalus*, *E. spinax* and *N. furzeri* allowed us i) to validate the lipofuscin and 3-NT accumulation as new cardiac aging markers in *S. microcephalus* heart; ii) to distinguish *S. microcephalus* features of cardiac aging associated to centenary lifespan versus adaptations associated with deep-sea habitat or phylogenetic lineage.

## 2 Materials and methods

### 2.1 Tissues sampling and processing

For paraffin embedding and Electron microscopy (EM): *S. microcephalus* heart samples were collected from six *S. microcephalus*, that were captured by long-line fishing in Greenland waters. (Supplmentary table 1). Hearts were dissected immediately post-mortem and 1cm x 1cm size piece of myocardium taken from myocardial surface until lumen, mid way down the ventricle between the apex and base, recovered in 4% paraformaldehyde (PFA), fixed for 48h, then PBS washed and dehydrated trough ethanol (EtOH) 25%, 50%, 70% series. For EM: compact and spongy myocardium tissues were sampled separately. *E. spinax* heart samples were collected from seven specimens classified as juveniles (Follesa & Carbonara 2019) (Supplementary table 2) captured by deep-water trawlers operating in the Ligurian Sea (Santa Margherita Ligure). Hearts were dissected and processed as described for *S. microcephalus. N. furzeri* heart samples were collected from three females and three males of MZM-0410 strain at 39 weeks of age. Animals were euthanized with Tricaine, MS-222, in accordance with the prescription of the European Union (Directive 2010/63/UE) and hearts dissected and fixed in 4% PFA overnight (O.N.). Samples were PBS washed and dehydrated through EtOH series. All paraffin samples were sectioned 6µm thin.

### 2.2 Masson’s trichrome staining

Masson’s trichrome staining (Bioptica) were performed on deparaffinized sections according routine procedures for light microscopy with the following modifications: nuclei were stained with Weigert’s Iron Hematoxylin Solution for 10 min and then glass slides H2O washed before proceeding with Masson’s trichrome staining.

### 2.3 Cardiac fibrosis quantification

For the fibrosis quantification we analyzed three animals for each species, two males and one female for *S. microcephalus* and two female and one male *for E. spinax*. For each animal three different field of Masson’s trichrome stainings were imaged in the spongy and compact myocardium layer in white light, all images were captured under identical illumination, exposure, and white balance settings to maintain consistency across samples. For each image, fibrosis was quantified based on selective segmentation of the fibrotic signal. Briefly, images were converted to the Hue–Saturation–Brightness (HSB) color space, and each channel was thresholded independently. Logical and combination of thresholded Hue, Saturation, and Brightness channels yielded a binary fibrosis mask representing collagen-positive areas. Nuclei were quantified from the same image field to normalize fibrosis to the numbers of nuclei. Original brightfield images were converted to 8-bit grayscale and thresholded to segment nuclei. Watershed separation was applied to resolve adjacent nuclei. For each image, the fibrosis burden was expressed as: Fibrosis Ratio = Fibrosis Area (pixels) /Number of Nuclei. Fibrosis ratios were further analyzed with GraphPad Prism (version 6.1, GraphPad Software, San Diego, CA, USA) for data analysis and visualization. Data were plotted as mean ± SEM for each animal, statistical comparisons between two groups were performed using an unpaired Student’s t-test.

### 2.4 Sudan Black B staining

Deparaffinized sections were stained in Sudan B black (Sigma Aldrich, Cat # S0395) 0.5% solution for 10 min RT, and washed three times in H_2_O 10 min.

### 2.5 Photobleaching

Deparaffinized slices were stained with Hoechst 33342 2µM 1:10000 for 5 min, washed three times in PBS. Images were acquired using an epifluorescence microscope (Nikon EclipseE600) equipped with a DS-Fi3 color camera (Nikon, Tokyo, Japan). Photobleaching was performed exposing the samples to continuous fluorescence blue-light illumination (DAPI filter) for 30 min.

### 2.6 Lipofuscin quantification

For lipofuscin quantification we analyzed three females for each species. For each animal three different fields were imaged within the spongy myocardium layer with an epifluorescence microscope in the DAPI channel. Fluorescence images were analyzed using Fiji (ImageJ, NIH, version 1.54p) as follow: nuclei were identified in the DAPI channel using a semi-automated macro workflow. Images were first converted to 8-bit grayscale, nuclei were segmented by automatic thresholding (Otsu method) followed by binarization and separation of touching objects using the *Watershed* algorithm. Nuclei were quantified using the *Analyze Particles* tool. All processing parameters were kept constant across samples. Nuclei masks were then generated and their fluorescence subtracted from the corresponding image. The resulting nuclei-subtracted image was then analyzed over the entire field and the mean gray value recorded. Mean gray values were further analyzed with GraphPad Prism (version 6.1, GraphPad Software, San Diego, CA, USA). Data were plotted as mean ± SEM for each animal, statistical comparisons between two groups were performed using an unpaired Student’s t-test. Masson’s trichrome, Sudan Black B staining, photobleaching and lipofuscin were imaged with an epifluorescence microscope (Nikon EclipseE600) equipped with a DS-Fi3 color camera (Nikon, Tokyo, Japan).

### 2.7 Electron Microscopy (EM)

Tissues were immediately excised post mortem, immersed in Karnovsky fixative (2.5% glutaraldehyde, 2% PFA in 0.1 M sodium cacodylate buffer, pH 7.2–7.4) (Pinali & Kitmitto 2014), cut into ∼1 mm³ cubes, and stored at 4 °C until further processing. Samples were post-fixed and stained en bloc following a protocol adapted for cardiomyocytes (Deerinck et al. 2010; Smith & Starborg 2019). After washing in 0.1 M sodium cacodylate buffer, samples were post-fixed in reduced osmium (2% osmium tetroxide, 1.5% potassium ferrocyanide) for 1 h, rinsed in double-distilled water (ddH₂O), incubated in filtered thiocarbohydrazide for 1 h at RT, and washed again in ddH₂O. They were subsequently treated with 1% osmium tetroxide for 1 h, washed, incubated in 1% uranyl acetate overnight at 4 °C, washed, and finally incubated in Walton’s lead aspartate at 60 °C for 30 min before a final rinse in ddH₂O. Dehydration was carried out through graded EtOH (30%, 50%, 70%, 90%, 100% and 100%) followed by pure acetone. Samples were infiltrated with increasing concentrations of TAAB 812 Hard Resin (TAAB Laboratories Equipment Ltd, UK) in acetone (25% resin in acetone, 50%, 75% up to 100 % resin) and polymerised at 60 °C for 24–48 h. Resin blocks were mounted on aluminium pins, trimmed to expose a 500 µm × 500 µm face, polished with a diamond knife, and gold-coated prior to imaging. Volumetric imaging was performed using an FEI Quanta 250 FEG scanning electron microscope (FEI Company, USA) equipped with a Gatan 3View serial block-face system (Gatan, Inc., USA). Image stacks were acquired at 3.8 kV, 0.45 Torr chamber pressure, and 3.5 μs/pixel dwell time, at a calibrated magnification of 12 nm/pixel. Serial sections of 50 nm were cut, producing image stacks of several hundred slices at 6,000 × 6,000 pixels. The resulting voxel dimensions were 12 nm (x, y) and 50 nm (z).

### 2.8 Immunohistochemistry (IHC)

Immunohistochemistry were carried out according to standard procedures. Primary antibodies were all incubated overnight at 4 °C according to the following dilutions: sarcomeric α-actinin (Sigma Cat# A7811) 1:400, 3-nitrotyrosine 3-NT (Invitrogen Cat# A-21285) 1:200, lysosomal Lamp1 (Abcam Cat# ab24170) 1:500. After the incubation sections were washed three times in PBS 1x and incubated 2 h with secondary antibodies 1:500 (Invitrogen polyclonal rabbit Alexa Fluor 488 Conjugated Cat# A-11008, RRID:AB_143165; Invitrogen polyclonal mouse Alexa Fluor 568, Cat# A-11004). Nuclei were stained with Hoechst 1:5000 5 min and sections mounted with fluoroshield mounting medium. IHC Imaging was performed using a confocal STELLARIS 8 (Leica) microscope and operated via LAS X software (Leica Microsystems, Wetzlar, Germany). High resolution Airyscan imaging was performed with confocal Zeiss LSM900 microscope (Carl Zeiss) equipped with Airyscan 2, and operated via ZEN 3.1 (blue edition) software.

## 3. Results

### 3.1 Cardiac fibrosis

We started our investigation by assessing the histology of the *S. microcephalus* ventricle. The ventricular morphology of *S. microcephalus* is consistent with that of low-activity elasmobranchs, characterized by a rigid outer pericardial layer predominantly composed of collagen (Figure S1), and a compact myocardial layer overlying a widespread spongy myocardium. Additionally, a well-developed coronary artery network supplying the entire ventricle is present (Figure S1). Masson’s trichrome staining revealed widespread fibrosis throughout the ventricular myocardium of *S. microcephalus*, affecting both the external compact layer and the internal spongy myocardium, belonging to the interstitial and perivascular fibrosis subtype (Fig. 1AF, Supplementary S2,A–C). We analyzed six *S. microcephalus* ranging 300–340 cm TL. This size corresponds roughly to 100–150 years and the time of sexual maturity in females (Nielsen et al. 2016). Collagen infiltration was observed in all six specimens and affected both myocardial sublayers and both sexes (Fig. 1A; B; D; E) (Fig. 1C; F). To exclude that extensive collagen deposition is a general trait associated with the deep-sea environment, we performed histological analysis of the ventricular myocardium of the deep-sea squaliform shark *Etmopterus spinax*. Masson’s trichrome staining of *E. spinax* ventricular wall revealed a canonical shark myocardial architecture, with an outer fibrotic pericardial layer underlied by compact and spongy myocardial layer. No evidence of cardiac fibrosis was detected in the ventricular wall of either females or males in a total of seven specimens (Fig. 1G–L). Quantification of fibrotic tissue in the two sharks, two males and one female for *S. microcephalus* and two female and one male for *E. spinax*, clearly showed the absence of cardiac fibrosis both in compact and spongy myocardium of *E. spinax*; while the *S. microcephalus* showed a massive fibrotic presence in both layer (Fig. 1R). To further determine if the massive cardiac interstitial fibrosis is a general phenotypic characteristic of aged fish, or it is specific of the extremely long-living *S. microcephalus*, we performed histological analysis of the ventricular myocardium of two males and one female at 39 weeks of age (i.e. median lifespan) of the short living teleost *N. furzeri*, a model where age-dependent cardiac changes are well described (Ahuja et al. 2019; Heid et al. 2017; Ma et al. 2025). We did not detect interstitial cardiac fibrosis (Fig. 1M–Q) but observed occasionally cardiac scars reminiscent of ischemic lesions (n = 3, two males and one female of the samples evaluated), a representative lesion is illustrated in Fig. 1O, yellow arrow. The presence of such scars highlights the capability of killifish myocardium to generate collagen rich fibrotic scars when focally injured, thus excluding that absence of interstitial fibrosis is due to inability to mount a fibrotic injury response.

**Figure 1:**
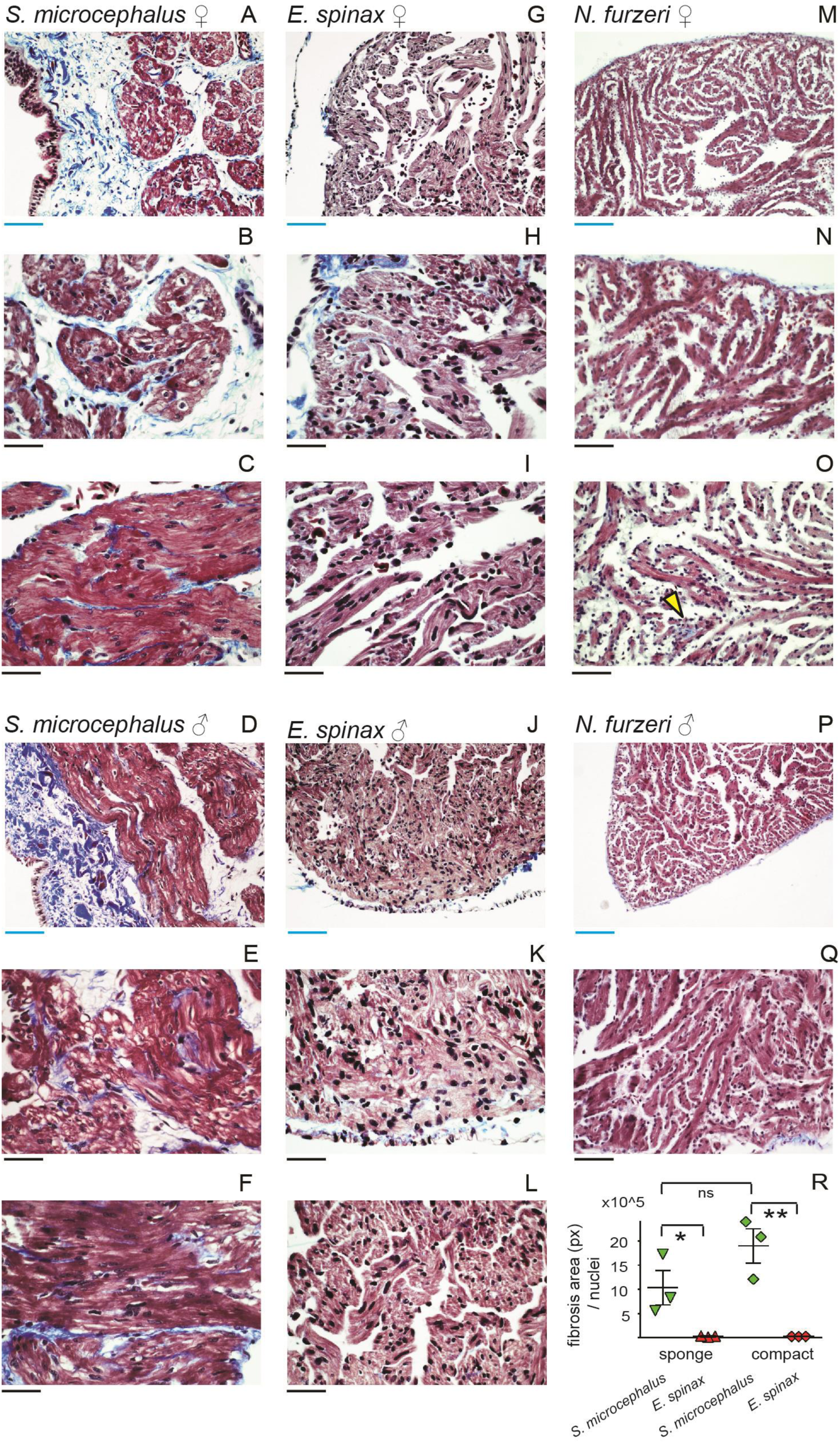
Histological evaluation and cardiac fibrosis of several fish species. A, B) *S. microcephalus* 341cm TL female, compact layer. C) *S. microcephalus* 341cm TL female, spongy layer. D, E) *S. microcephalus* 335cm TL male. F) *S. microcephalus* 335 TL male, spongy layer. G, H) *E. spinax* 300mm TL female, compact layer. I) *E. spinax* 300mm TL female, spongy layer. J, K) *E. spinax* 260mm TL male, compact layer. L) *E. spinax* 260mm TL male, spongy layer. M–O) *N. furzeri* ventricular myocardium, 39 weeks old female. P,Q) *N. furzeri* ventricular myocardium, 39 weeks old male. Blue scalebar: 100µm, black scalebar: 50 µm. yellow arrow: ischemic collagen scar. R) Quantification of cardiac fibrosis in sharks.

### 3.2 Lipofuscin accumulation

To explore whether extreme longevity of *S. microcephalus* correlates with expression of cellular aging biomarkers, we analyzed lipofuscin deposition in the ventricular wall of *S. microcephalus*, *E. spinax* and aged *N. furzeri*. Sudan Black B staining of ventricular myocardial sections revealed massive and diffuse lipofuscin accumulation filling the entire cardiomyocyte in *S. microcephalus*, both in the compact and spongy myocardial layers, in both females and males (Fig. 2A–F). In contrast, *E. spinax* exhibited limited Sudan Black–positive areas, primarily restricted to localized regions of the compact myocardium (Fig. 2G–L), and with markedly lower staining intensity as compared to *S. microcephalus*. Interestingly, aged *N. furzeri* showed abundant lipofuscin granules, although these were detectable only at higher magnification and did not fill the sarcoplasm of the cardiomyocytes. Unlike in sharks, the pigment was localized mainly outside the cardiomyocytes, dispersed within the trabecular interspaces (Fig. 2M–P, black arrows). To confirm lipofuscin accumulation, we exploited its intrinsic autofluorescence, which typically emits across the green-to-red spectrum (approximately 540–640 nm). Comparative autofluorescence analysis of myocardial sections from *S. microcephalus*, *E. spinax*, and aged *N. furzeri* revealed the characteristic golden-yellow autofluorescence of lipofuscin granules under blue-light excitation (DAPI filter). In *S. microcephalus*, prominent lipofuscin accumulation was observed within cardiomyocytes of both the compact and spongy myocardial layers, in specimens of both sexes (Fig. 3A–D). By contrast, no apparent lipofuscin autofluorescence was detected in *E. spinax* (Fig. 3E–H) even though autofluorescence of erythrocytes was readily visible in both species (Fig. 3D, F; red arrows). As expected, lipofuscin quantification between *S. microcephalus* and *E. spinax* highlighted a statistically significant difference in lipofuscin content between the two shark species (Fig. 3K). as previously assumed, aged *N. furzeri* displayed numerous golden autofluorescent granules, consistent with lipofuscin, located primarily within the trabecular interspaces rather than inside cardiomyocytes (Fig. 3I–J). Given the Sudan Black B well-documented ability to quench lipofuscin’s intrinsic autofluorescence, we extended our analysis by imaging *S. microcephalus* and aged *N. furzeri* myocardial sections stained with Sudan Black B under fluorescence light across three major spectral channels: blue (DAPI filter), green (GFP/FITC filter), and red (TRITC/CY3 filter). As anticipated, *S. microcephalus* samples exhibited marked attenuation of golden autofluorescence in the blue spectrum (Fig. 4A), confirming Sudan Black B-mediated quenching, while dense deposits of Sudan Black B-positive lipofuscin remained visible (Fig. 4B–C) under white light. To further validate that the observed autofluorescence originated from lipofuscin, we conducted a photobleaching experiment, capitalizing on the well-documented resistance of lipofuscin to photobleaching (Sun & Chakrabartty 2016; Davies et al. 2001). Myocardial sections were exposed to continuous blue-light illumination (DAPI filter) for 30 min. Following exposure, we observed marked photobleaching of the nuclear Hoechst staining (Fig. 4D–E), confirming effective light-induced degradation of canonical fluorophores. By contrast, the strong golden autofluorescent signal attributed to lipofuscin persisted within cardiomyocytes and remained clearly visible post-bleaching (Fig. 4F), consistent with lipofuscin’s photostability. As expected, in the aged heart of *N. furzeri*, photobleaching revealed an abundance of persistent golden autofluorescent granules localized predominantly in the trabecular interspaces, further supporting their identification as lipofuscin (Fig. 4G). Moreover, the *N. furzeri* accumulation of lipofuscin outside the cardiomyocytes is in line of what was already observed in mice hearts (Wang et al. 2022). These granules remained detectable in the far-red fluorescence channel of a confocal microscope, further supporting their identification based on lipofuscin’s broad emission spectrum and resistance to photobleaching (Fig. 6S, V).

**Figure 2:**
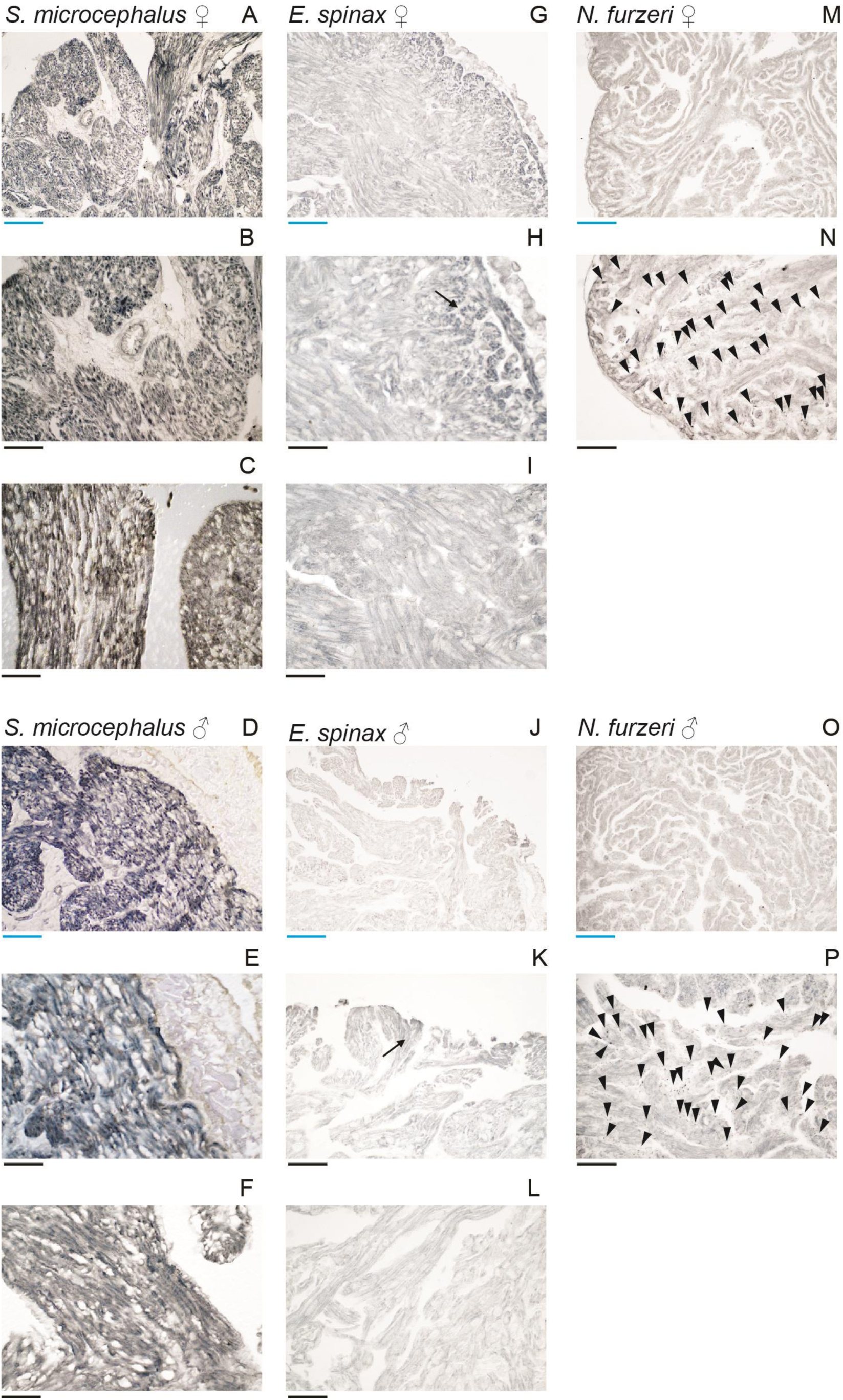
Sudan Black B staining of ventricular myocardium. A, B) *S. microcephalus* 341cm TL female, compact layer. *S. microcephalus* female, compact layer. C) *S. microcephalus* 341cm TL female, spongy layer. D, E) *S. microcephalus* 310cm TL male. F) *S. microcephalus* 310cm TL male, spongy layer. G, H) *E. spinax* 300mm TL female, compact layer, black arrow. I) *E. spinax* 300mm TL female, spongy layer. J, K) *E. spinax* 260mm TL male, compact layer, black arrow. L) *E. spinax* 260mm TL male, spongy layer. M-N) *N. furzeri* 39weeks old ventricular myocardium, female, lipofuscin granules, black triangles. O–P) *N. furzeri* 39weeks old ventricular myocardium 39 weeks old male, lipofuscin granules, black triangles. Blue scalebar: 100µm, black scalebar: 50 µm

**Figure 3:**
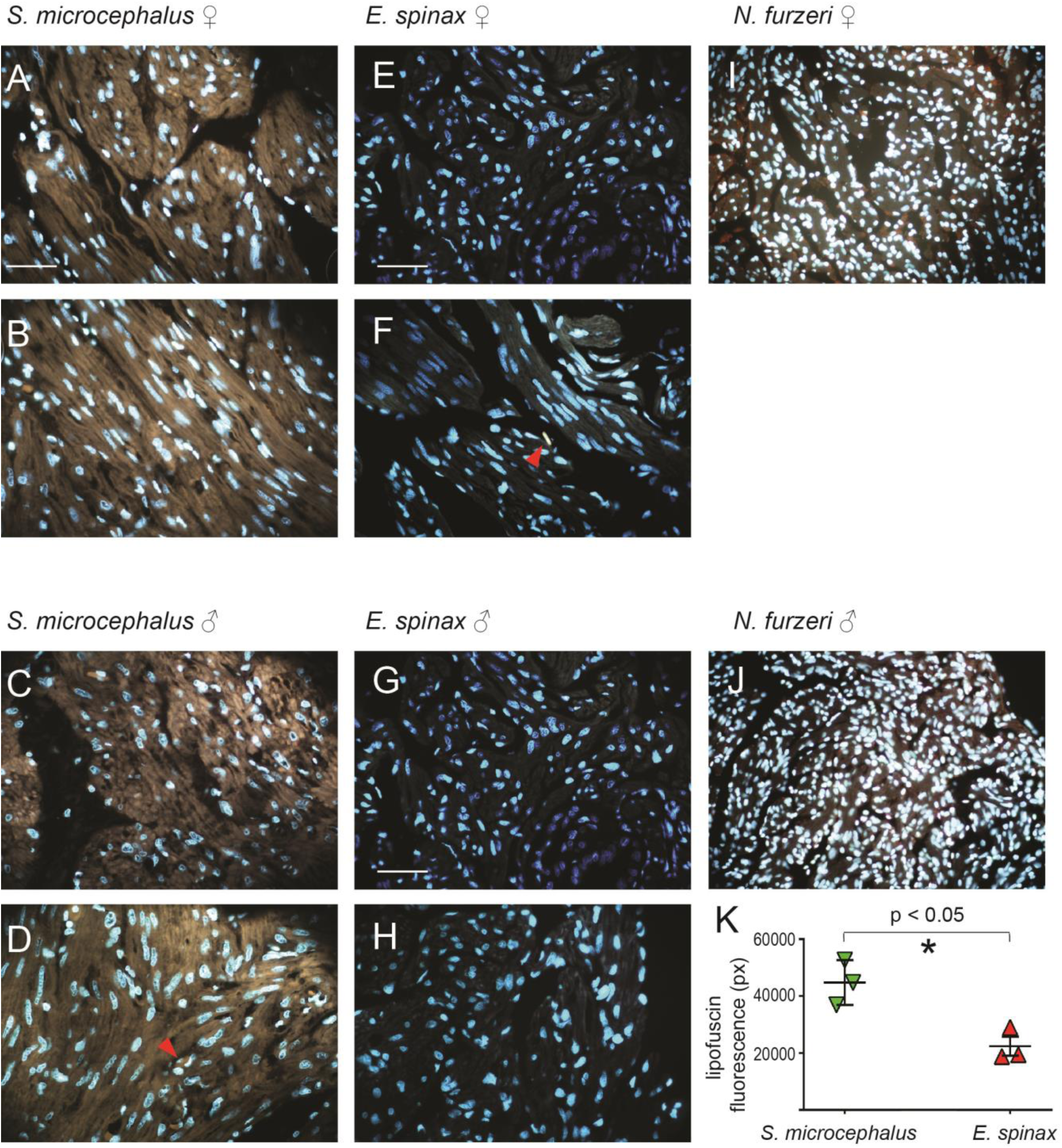
Myocardium autofluorescence highlighting lipofuscin. A) *S. microcephalus* 325cm TL female, myocardial compact layer. B) *S. microcephalus* 325cm TL female, myocardial spongy layer. C) *S. microcephalus* 310cm TL male, myocardial compact layer. D) *S. microcephalus* 310cm TL male, myocardial spongy layer. E) *E. spinax* 300mm TL female, myocardial compact layer. F) *E. spinax* 300mm TL female, myocardial spongy layer. G) *E. spinax* 260mm TL male, myocardial compact layer. H) *E. spinax* 260mm TL male, spongy layer. I) *N. furzeri* ventricular myocardium 39 weeks old female. J) *N. furzeri* 39 weeks old ventricular myocardium, male. K) lipofuscin quantification in *S. microcephalus* and *E. spinax*.

**Figure 4:**
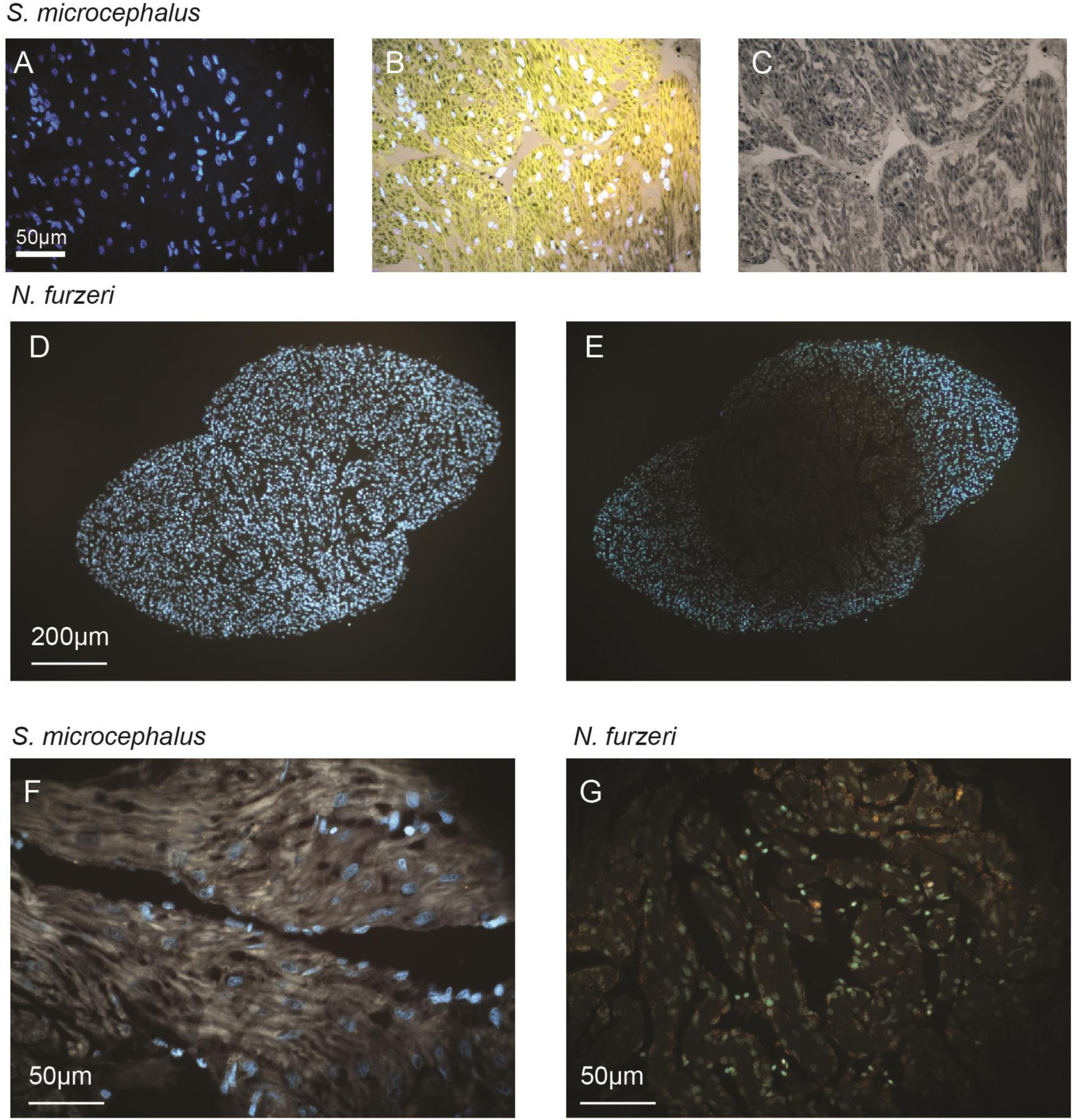
lipofuscin in the ventricular myocardium. A) Sudan Black B stained ventricular myocardium of *S. microcephalus*, acquired with DAPI-filter fluorescent light showed quenched lipofuscin autofluorescence, with Hoechst in blue. B) RGB autofluorescence acquired in the TRITC/CY3, GFP/FITC and DAPI filters, respectively and merged on the white light acquisition. Sudan Black B granules quenched the RGB yellow-resulting basal autofluorescence. Merge. C) White light acquisition. Representative images of coronal section of *N. furzeri* ventricle before D) and E) after photobleaching. F) Photobleached acquisition of a spongy ventricular *S. microcephalus* section shows clear golden autofluorecence. G) Photobleached acquisition of *N. furzeri* ventricle shows clear golden autofluorecence granules. A, B, D–G cyan: Hoechst acquired with DAPI-filter. A, D–G golden: lipofuscin autofluorescence acquired with DAPI-filter.

### 3.3 Ultrastructural analysis

To further investigate the subcellular nature of lipofuscin accumulation, we performed electron microscopy (EM) of the *S. microcephalus* myocardial compact layer. Lipofuscin is a by-product of lipid peroxidation that accumulates within lysosomes as an indigestible amorphous aggregate. Damaged mitochondria, due to oxidative stress, are one of the main sources of the oxidized material that becomes lipofuscin and accumulate into autophagosomes (Terman et al. 2004; König et al. 2017; Lu et al. 2020). We analyzed three *S. microcephalus* specimens (one male and two females) ranging from 294 to 434 cm TL. In all specimens we observed a massive accumulation of electrondense, lipofuscin-packed autophagosomes (Fig. 5A–E white arrows), together with numerous lysosomes filled with electrondense material (Fig. 5A–E, red arrows). These were embedded within a dense network of mitochondria. In addition, we identified degenerating mitochondria apparently in the process of being phagocytosed and incorporated into autophagosomes (Fig. 5B–E, yellow arrows). The autophagosomes exhibited variable morphology and size, ranging from mitochondrion-like profiles (Fig.5B, E white arrows) to structures several micrometers in size (Fig. 5A, C, D, white arrows), supporting the hypothesis that lipofuscin can accumulate within *S. microcephalus* cardiomyocytes in very large amounts without compromising their viability or functionality. To further confirm ultrastructural data we perform high resolution Airyscan imaging on *S. microcephalus* compact and spongy myocardial layer of a 341cm TL female and 335cm TL male aiming to image lysosomal and lipofuscin granules. Single plane acquisitions showed i) myocardial lipofuscin distribution consistent with Sudan Black B staining in both the compact and spongy myocardium of the two animals analyzed (Fig. 4B–C; fig 5F–U); ii) the granular-shape nature of the lipofuscin structures, consistent with EM data (Fig. 5F, G, I, J, K, M, N, O, Q, R, S, U) iii) the general colocalization of lysosomal Lamp1 with the lipofuscin granules (Fig. 5G–I, K–M, O–Q, S–U). Colocalization is variable in Lamp1 fluorescence intensity (Fig.5F–U, single examples in G, K, O, S: orange arrows), and some lipofuscin structures are Lamp1 negative (Fig.5 F–U, single examples in G, K, O, S: violet arrows), while lysosomes lipofuscin-negative are clearly distinguishable all around the myocardium (Fig.5 G, K, O, S), indicating the specificity and the robustness of the IHC staining. Together, the EM and IHC findings support the hypothesis that the large autophagosomes represent fusion products of lysosomes with degenerating mitochondria.

**Figure 5:**
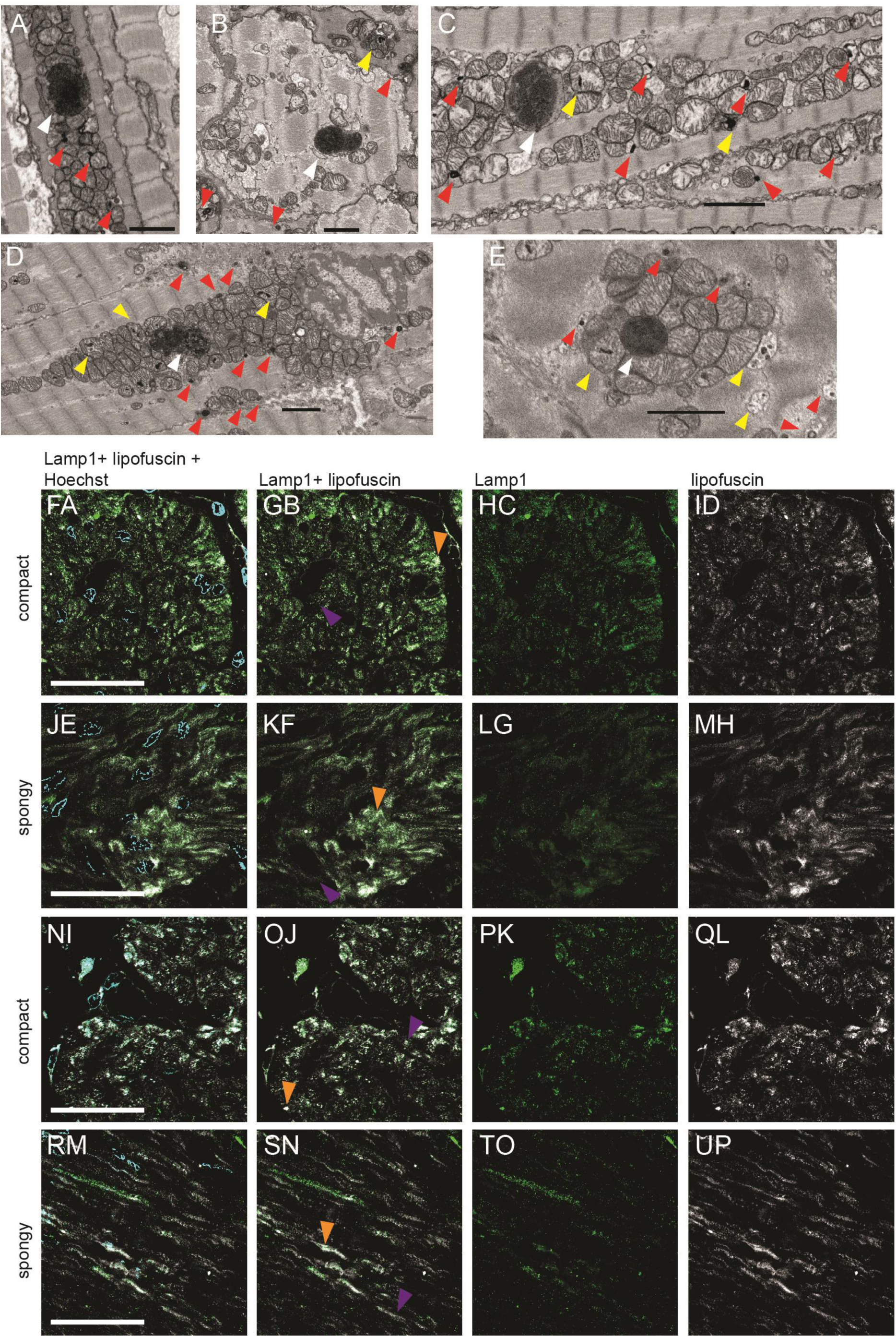
EM of *S. microcephalus* showing lipofuscin accumulation. A–D) EM of a 294cm male. E, F) EM of a 310cm female. G, H) EM of a 434cm female. White arrow: autophagosomes. Red arrow: lysosomes. Yellow arrow: dying mitochondria in the process to be incorporated inside autophagosomes. Scalebar 2µm.

**Figure 6:**
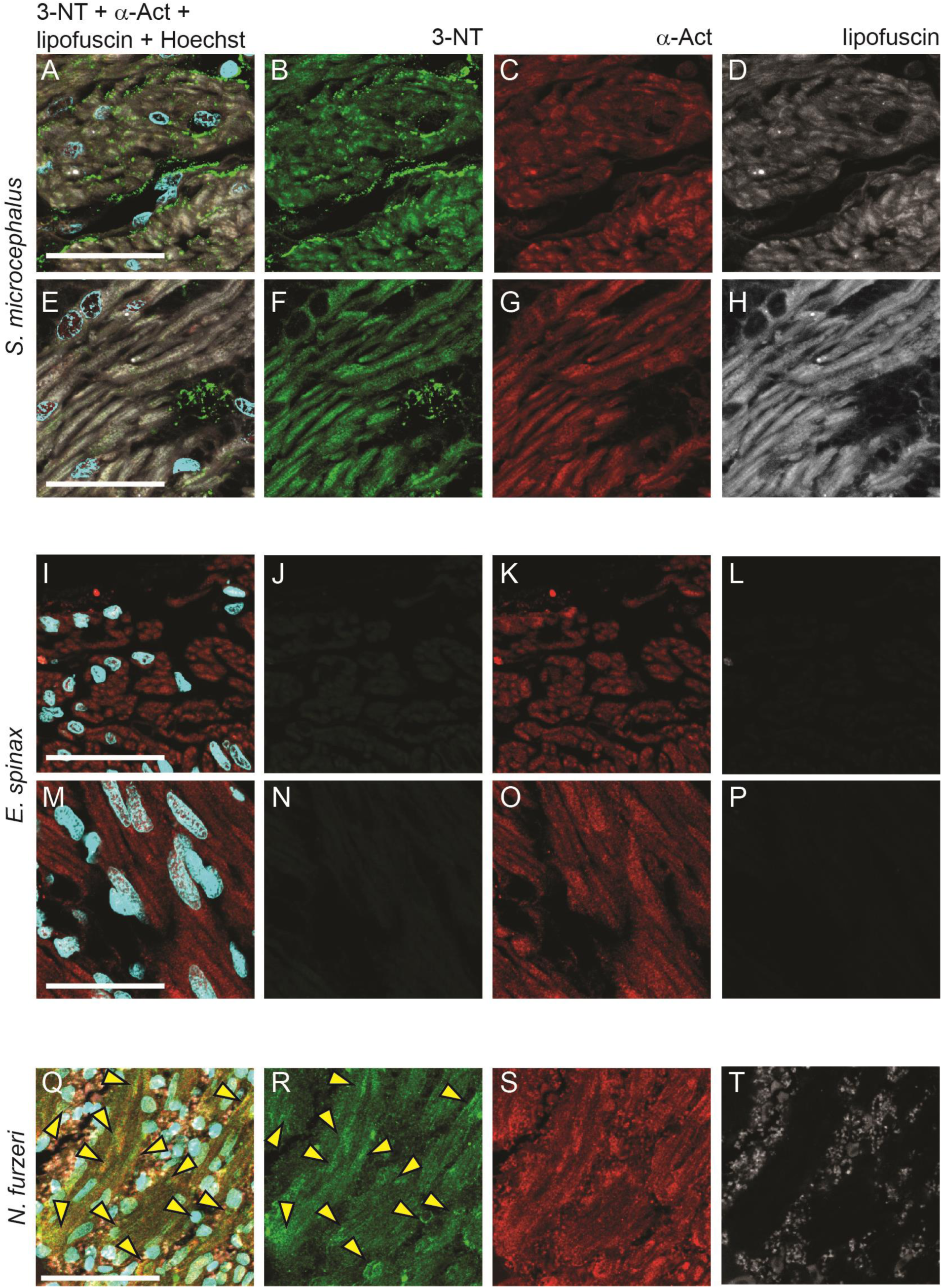
IHC of *S. microcephalus* showing lysosomes and lipofuscin accumulation acquired with Airyscan. A–H) IHC of 341cm TL female myocardium. I–P) IHC of 335cm TL male myocardium. Yellow arrows: single examples of lysosmal Lamp1 and lipofuscin colocalization. Red arrows: single examples of lipofuscin only granules. Cyan: Hoechst. Green: Lamp1. White: lipofuscin. Scalebar 50µm.

### 3.4 Nirotyrosine accumulation

3-Nitrotyrosine (3-NT), a stable product of tyrosine nitration by reactive nitrogen species, is widely recognized as a hallmark of cardiac aging and a reliable biomarker of oxidative and nitrosative stress. Its accumulation has been reported in cardiac tissue across several species, including the *N. furzeri* (Heid et al. 2017). We investigated 3-NT presence in the ventricular myocardium of S. microcephalus, *E. spinax* and aged *N. furzeri*. Strong immunoreactivity for 3-NT (Fig. 6A–H) was detected in both male and female specimens in the ventricular myocardium of *S. microcephalus*, with prominent staining observed in both the compact (Fig. 6A, B) and spongy myocardial layers (Fig.6E, F). Autofluorescence imaging in the far-red channel confirmed the presence of lipofuscin within cardiomyocytes, consistent with earlier observations (Fig.6A, D, E, H). In contrast, *E. spinax* myocardial tissue showed no detectable 3-NT signal (Fig. 6I–R) and lacked lipofuscin-associated autofluorescence (Fig. 6I, L, M, P). Interestingly, nuclei within the spongy myocardium of *E. spinax* appeared enlarged as compared to those of the other species analyzed (Fig. 1H, I, K, L; Fig. 3B, D, F, H; Fig. 6M), suggesting a species-specific morphological trait. Notably, in *S. microcephalus*, 3-NT deposition was particularly abundant in interstitial regions, whereas its presence within cardiomyocytes was comparable to that observed in aged *N. furzeri* (Fig. 6Q–T) and already reported in literature (Heid et al. 2017).

## 4. Discussion

Our study provides the first histological and molecular analysis of cardiac aging in the Greenland shark (*Somniosus microcephalus*), a species notable for its extraordinary lifespan exceeding four centuries. Human centenarians show three different phenotypes: some show no (escapers) or delayed (escapers) onset of age-associated diseases, others maintain physiological functions despite the presence of pathological conditions (survivors). These conditions are exemplary of resistance vs. resilience. Resistance is the capacity to oppose a perturbation, resilience can be defined as the capacity of a system to return to a physiological state after a perturbation. Despite harboring hallmark features of cardiac aging such as interstitial and perivascular fibrosis, lipofuscin accumulation, and 3-NT deposition, *S. microcephalus* individuals appeared phenotypically healthy. More strikingly, these molecular and cellular aging traits were observed in relatively young individuals of 300cm to 340cm, far from the 500 cm maximum length. This unique decoupling of aging markers from functional decline underscores resilience as a key mechanism enabling extreme longevity and highlights the Greenland shark as a unique model for studying resilience to tissue aging.

In our analysis we observed extensive interstitial and perivascular fibrosis in the ventricles of *S. microcephalus*, spanning both the compact and spongy myocardial layers. Cardiac fibrosis is a well-documented hallmark of aging across vertebrate taxa, including humans, rodents, and teleosts, and is generally associated with maladaptive remodeling, reduced diastolic compliance, and increased risk of arrhythmogenesis (Schulman et al. 1992; Gazoti Debessa et al. 2001; LEVY et al. 1988; Chen et al. 2022; Kane et al. 2021; Dai & Rabinovitch 2009; Keen et al. 2015). Age-associated cardiac fibrosis is typically interstitial and diffuse, rather than focal or replacement in nature. Histologically, it is marked by an accumulation of collagen type I and type III, predominantly in the interstitial and perivascular regions (Biernacka & Frangogiannis 2011). Surprisingly, this fibrotic signature was not associated with any visible signs of pathology or dysfunction in the adult examined individuals of *S. microcephalus*. These findings are consistent with recent reports of non-pathological, age-related cardiac fibrosis in mice, described as a hallmark of cardiac aging (Basilicata et al. n.d.). Notably, neither the deep-sea elasmobranch *E. spinax* nor the aged teleost *N. furzeri* displayed comparable fibrotic remodeling. The absence of fibrosis in *E. spinax*, a species that inhabits comparable deep-sea environments but exhibits a markedly shorter lifespan, suggests that the extensive fibrosis observed in *S. microcephalus* is not simply an adaptive response to hydrostatic pressure. On the other hand, the absence of interstitial fibrosis in the aged heart of *N. furzeri* support the hypothesis that aging processes alone are not sufficient to develop collagen deposition and interstitial fibrosis formation. Rather, our observations support the hypothesis that *S. microcephalus* has evolved an exceptional resilience to age-related cardiac fibrosis, maintaining physiological function despite extensive extracellular matrix remodeling. In the Mediterranean Sea, the deep-water temperature structure is comparatively stable, with a persistent thermocline and typical temperatures of approximately 12.8–13.2 °C below ∼200 m depth (Houpert et al. 2015; Cardin et al. 2015). In several teleost freshwater species, including, common carp *Cyprinus carpio* (Goolish 1987), channel catfish *Ictalurus punctatusi* and green sunfish *Lepomis cyanellus* (Kent et al. 1988) chronic cold acclimation has been shown to induce cardiac ventricular hypertrophy. In anadromous salmonids such as rainbow trout *Oncorhynchus mykiss* and atlantic salmon *Salmo salar*, cold-induced cardiac fibrosis has also been documented (Farrell et al. 1988; Keen et al. 2015). Such cold-associated cardiac remodeling in teleosts is generally interpreted as an adaptive response, producing a hypertrophied ventricle capable of generating the necessary contractile force to maintain hemodynamic function at low temperatures while limiting overstretch under conditions of increased blood viscosity in species which can not reduce physical activity during seasonal temperature changes. For reference, *O. mykiss* typically exhibits arterial blood pressures of ∼3–6 kPa and resting heart rates of 20–50 bpm (Le Mével et al. 2002), whereas *S. salar* shows values of ∼4–7 kPa and ∼56 bpm (Perry et al. 1999). Blood pressure in Greenland sharks has never been directly measured, still it has been estimated by analyzing the relative amounts of elastin and collagen in the wall of the ventral aorta, together with measuring its compliance characteristics over a range of pressures (Shadwick et al. 2018). These data suggest that the Greenland shark’s average blood pressure is approximately 2.3–2.8 kPa (Shadwick et al. 2018), much lower than other sharks. For example the *Scyliorhinus canicula* (commonly known as small spotted catsharks) is an animal with a relatively low activity and metabolic rate, still possess a blood pressure of 3–5 kPa (Taylor et al. 2003). Other sharks (e.g. lamnid sharks) have even higher metabolism and blood pressure, over >7 kPa; (Lai et al. 1997). Such comparatively low blood pressure in *S. microcephalus* may reduce the selective pressure for cold-induced ventricular stiffening or fibrosis. To date, cold-induced cardiac hypertrophy or fibrosis has not been documented in Elasmobranchs, suggesting that this form of thermal cardiac remodeling may represent an evolutionary adaptation specific to teleost fishes, particularly those inhabiting freshwater or anadromous environments. Supporting this hypotesis, such collagen deposition due to cold acclimation was demonstrated to be reversible in rainbow trout, in which warm acclimation mimicking seasonal cycles reduces the collagen content (Johnston & Gillis 2022). Nevertheless, the possibility that, at least in part, cold exposure may contributes to the pronounced fibrotic response observed in *S. microcephalus* cannot be entirely excluded. Previous studies on the cardiocirculatory system of *S. microcephalus* have shown that it possesses a ventral aorta with a relatively low abundance of elastic fibers, which are loosely organized. This structural arrangement contributes to high arterial compliance at relatively low blood pressure (Shadwick et al. 2018). We speculate that age-related cardiac fibrosis in *S. microcephalus* may be counterbalanced by the elastic properties of the aorta, whose high compliance at low pressure helps preserve healthy cardiac function throughout aging.

Lipofuscin accumulation is widely regarded as a robust indicator of cellular aging, reflecting the progressive buildup of undegradable, cross-linked oxidized proteins and lipids within lysosomes of postmitotic cells (Terman & Brunk 1998; Brunk & Terman 2002; Terman & Brunk 2006). In this study, we report massive intracardiomyocyte deposition of lipofuscin in *S. microcephalus*, visualized using Sudan Black B staining, intrinsic autofluorescence, and photobleaching resistance. The signal was consistent across sexes and myocardial layers, while it is absent in the *E spinax*, suggesting that this is a generalized and pervasive feature in adult individuals of *S. microcephalus* species. Importantly, the persistent golden autofluorescence of lipofuscin in *S. microcephalus* and *N. furzeri* after photobleaching confirms its identity and highlights the robustness of the staining methods employed. Recently, metabolomic and proteomic analyses of mouse hearts revealed elevated ROS levels in aged mice accompanied by a concomitant decline in antioxidant defenses, while lipidomic profiling showed a marked accumulation of lipids that may serve as precursors for lipofuscin formation (Basilicata et al. n.d.). The extensive presence of lipofuscin-loaded autophagosomes and lysosomes in *S. microcephalus* cardiomyocytes evidenced by EM data, suggests a remarkable tolerance to oxidative stress and impaired organelle turnover in the cardiac tissue. The accumulation of lipofuscin derived from damaged mitochondria indicates that, despite ongoing oxidative processes, these cells can maintain viability without activating excessive autophagy or undergoing apoptosis. This resilience may reflect an adaptive mechanism allowing long-lived species such as *S. microcephalus* to preserve cardiac function over extended lifespans.

The extent to which lipofuscin manifest in the heart of *S. microcephalus*, and whether such aging marker deposition differs from that observed in shorter-lived vertebrates, had not been previously explored. The confirmed accumulation of lipofuscin in *S. microcephalus* hearts underscores its significance as an aging marker and provides a valuable tool for investigating the biology of this exceptionally long-lived vertebrate. We did not have access to older individuals close to the species maximum size and it remains unknown whether lipofuscin accumulation and fibrosis plateaus in adult life or progress leading to an even starker phenotypes. Our analysis also revealed substantial deposition of 3-nitrotyrosine (3-NT) in the ventricular myocardium of *S. microcephalus*. As a well-established indicator of oxidative and nitrosative stress, 3-NT has been widely used to assess age-related redox imbalance in various tissues, including the heart. Age-associated increases in 3-NT have been documented in the cardiac tissue of several species, including the *N. furzeri* (Heid et al. 2017). In the present study we were able to document the 3-NT in the myocardial tissue of *S. microcephalus*, while it proved to be absent in *E. spinax*. The spatial distribution of 3-NT immunoreactivity in *S. microcephalus* cardiomyocytes mirrored the pattern seen in aged *N. furzeri* heart, with further evident 3-NT deposits in the interstitial space of both compact and spongy layers. The oxidative stress theory of aging posits that longevity is achieved through reduced production of reactive oxygen species (ROS) or enhanced capacity to mitigate their effects. Yet, the case of *S. microcephalus* suggests an alternative model: that longevity may instead rely on exceptional resilience to chronic oxidative damage rather than its outright prevention. Supporting this notion, previous studies have reported high levels of oxidative status in the muscle of *S. microcephalus*, despite its remarkably long lifespan (Costantini et al. 2017) and mirrors the observation that naked mole rats show very high levels of lipid peroxidation as compared to mice (Andziak et al. 2006). Perhaps the most striking finding of this study is that, despite pronounced molecular and histological signatures of aging, *S. microcephalus* individuals appeared outwardly healthy and physiologically uncompromised at capture. This contrasts with mammalian models, where such features are typically associated with declining cardiac performance, increased arrhythmic risk, and mortality. The preservation of function despite accumulation of canonical aging markers, which are not detrimental for the century-living *S. microcephalus*, would suggest that the species tolerates its aged heart in a way that preserves function. All together our findings raise the possibility that despite observable structural changes, *S. microcephalus* maintains effective cardiac function via compensatory mechanisms and extremely high resilience. This comparative approach offers a promising avenue to further uncover molecular and histological features that contribute to the remarkable physiological resilience of *S. microcephalus*, allowing its centenary long live. All together our data contribute foundational insight into how one of Earth’s longest-lived vertebrates manages cellular and tissue aging in a vital organ. The *S. microcephalus* may illustrate a rare instance in which cumulative aging markers, such as fibrosis, lipofuscin and 3-NT, are tolerated, or perhaps compensated, without compromising cardiac function, offering a novel paradigm for understanding longevity in vertebrates. These findings may also inform translational approaches to mitigate age-related cardiac decline in humans.

## CRediT Author Contributions

**Elena Chiavacci**: Investigation, Methodology, Validation, Conceptualization, Formal analysis, Visualization, Writing – original draft, Writing – review & editing. **Kirstine Fleng Steffensen, Emanuele Astoricchio, Amalie Bech Poulsen, Daniel Brayson**: Resources. **Pierre Delaroche**: Investigation, Methodology, Validation, Writing – review & editing. **Fulvio Garibaldi, Luca Lanteri, Giovanni Roppo Valente, Federico Vignati**: Resources, Formal analysis, Validation, Writing – review & editing. **Christian Pinali**: Validation, Writing – review & editing. **John Fleng Steffensen, Holly Shiels**: Resources, Methodology, Validation, Writing – review & editing. **Eva Terzibasi Tozzini**: Methodology, Funding acquisition. **Alessandro Cellerino**: Methodology, Conceptualization, Project administration, Supervision, Funding acquisition, Visualization, Writing – review & editing

## Authorship statement

All authors have read and agreed to the published version of the manuscript.

## Acknowledgments

We are grateful to Dr. Sabine Matz and Dr. Cinzia Caterino for the support in the histological preparation of *N. furzeri* specimens. We also thank the staff of The University of Manchester Electron Microscopy Core Facility (RRID:SCR_021147; Faculty of Biology, Medicine and Health) for their assistance, particularly David Smith (former technician), whose support and expertise in SBF-SEM were instrumental in collecting these data. This work was supported by the Italian Ministry of University and Research (MUR) [Funder ID: 10.13039/501100003407] under PRIN 2022—SHARKAGE (Grant number CUP: E53D23007660001) and the PNRR project “THE—Tuscany Health Ecosystem—Spoke 1 and Spoke 8.

## Ethics Statement

Sampling in Greenland was carried out following laws and regulations and with authorization from the Government of Greenland (Ministry of Fisheries, Hunting & Agriculture, permit Nr. 2020-26794 and 2023-6108 and document number 565466, 935119, 20179208, C-17-129, C-15-17, and C-13-16). Specimens of *E. spinax* were bycatch of commercial fisheries. *N. fuzeri* were laboratory-bred as described by Naumann et al. (Naumann et al. 2024). All experiments were performed using the killifish strain MZM-0410 in accordance with relevant guidelines and regulations. Fish were bred and kept in the fish facility of the Leibniz Institute on Aging – Fritz Lipmann Institute according to §11 of the German Animal Welfare Act under license number J-003798. Sacrifice and organ harvesting were performed according to §4(3) of the German Animal Welfare Act.

## Figure legends

**Figure 7:** IHC showing 3-NT distribution in the ventricular myocardium. A–D) *S. microcephalus* 341cm TL female compact layer. E–H) *S. microcephalus* 341cm TL female spongy layer. I–L) *E. spinax* female 300mm TL compact layer. M–P) *E. spinax* female 300mm TL spongy layer. Q–T) *N. furzeri* 39w old female cardiac ventricle myocardium. Yellow arrows: 3-NT accumulation. Scalebar 50µm.

