## Supplementary for "Resilience to cardiac aging in Greenland shark *Somniosus microcephalus*"

#### Supplementary figures

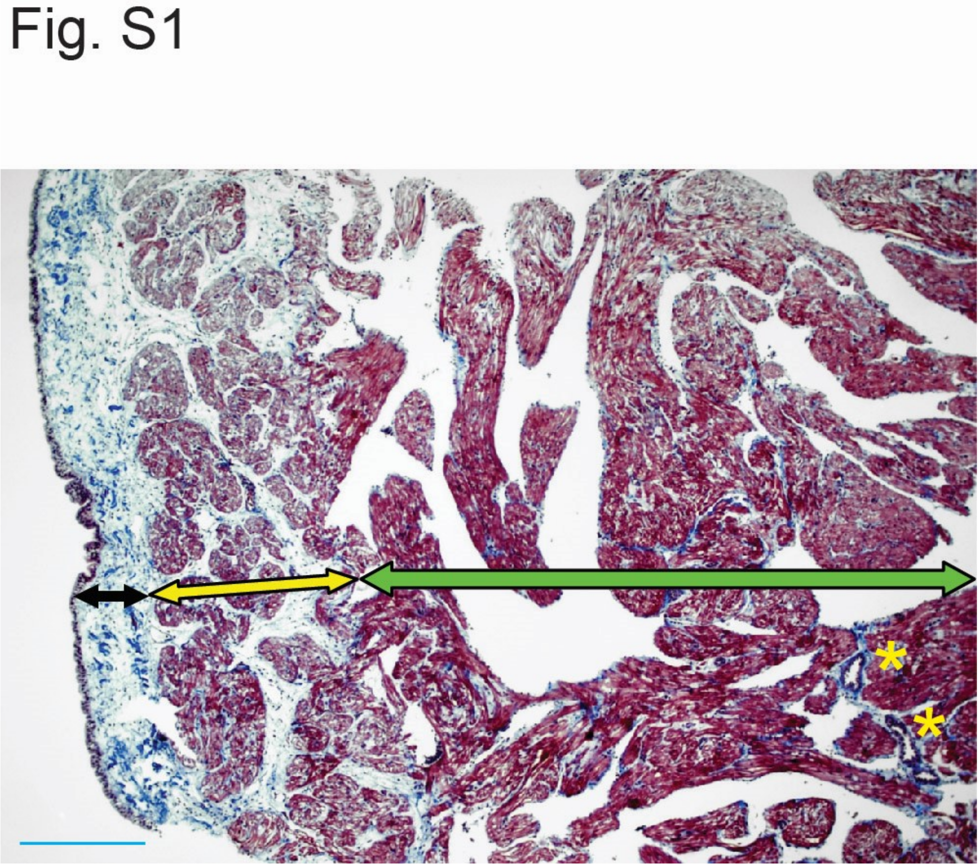


**Figure S1:** Representative of *S. microcephalus* ventricular wall. Black arrow: pericardial fibrotic capsule. Yellow arrow: compact myocardial layer. Green arrow: spongy myocardial layer. Yellow asterisks: coronary vessels. Blue scalebar: 500 µm.

Fig. S2

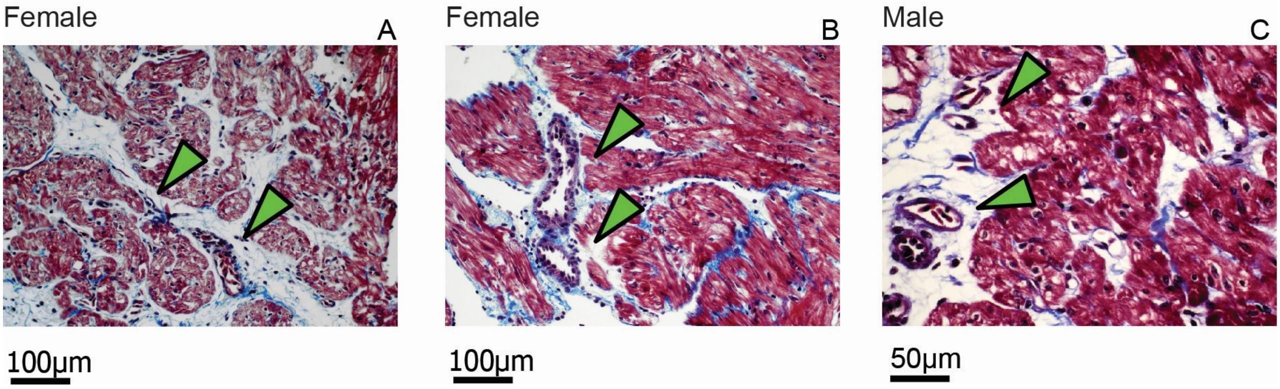


**Figure S2:** Representatives of *S. microcephalus* coronary cardiac fibrosis, green arrows: A) female 341cm; B) female 341cm; C) male 310cm.

Fig. S3

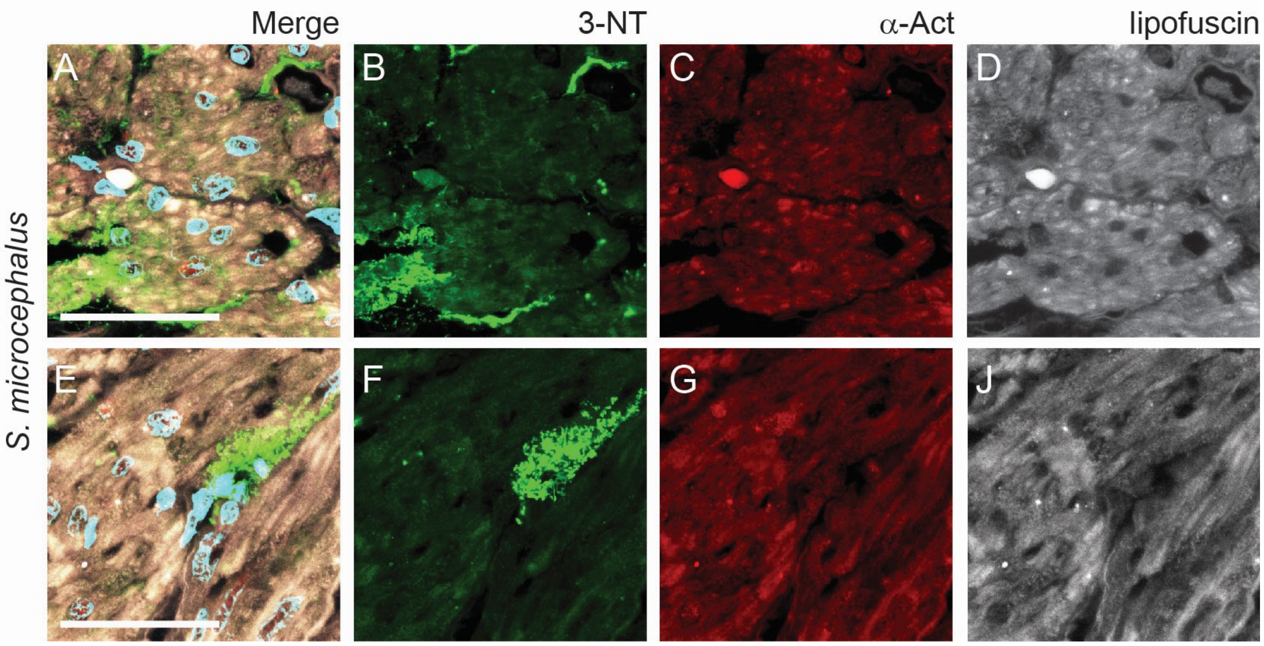


**Figure S3:** immunohistochemical 3-NT in the ventricular myocardium of a male *S. microcephalus* A) Compact layer, merge. B) Compact layer, 3-NT. C) Compact layer α-Actinin. D) Compact layer autofluorescent lipofuscin. E) Spongy layer merge. F) Spongy layer 3-NT. G) Spongy layer α-Actinin. J) Spongy layer autofluorescent lipofuscin. Scalebar, 50 μm.

### Supplementary tables

#### Supplementary table 1

| Samples | Sex | TL (cm) | Tissue processing |
|---|---|---|---|
| 1 | F | 303 | Paraffin |
| 2 | F | 325 | Paraffin |
| 3 | M | 335 | Paraffin |
| 4 | F | 341 | Paraffin |
| 5 | M | 310 | Paraffin |
| 6 | F | 330 | Paraffin |
| EM1 | M | 294 | EM |
| EM2 | F | 310 | EM |
| EM3 | F | 434 | EM |

Supplementary table 1: *S: microcephalus* parametres. F: female, M: male. EM: Electron Microscopy.

#### Supplementary table 2

| Samples | Sex | TL (mm) | Age ( years estimated) |
|---|---|---|---|
| 1 | F | 300 | 5.8 |
| 2 | F | 300 | 5.8 |
| 3 | F | 270 | 4.6 |
| 4 | F | 265 | 4.5 |
| 5 | F | 280 | 5.0 |
| 6 | F | 275 | 4.8 |
| 7 | M | 260 | 4.3 |

Supplementary table 2: *E: spinax* parametres. F: female, M: male.
